# Defining the impact of flavivirus envelope protein glycosylation on sensitivity to broadly neutralizing antibodies

**DOI:** 10.1101/2023.06.20.545776

**Authors:** Maya Contreras, Jackson B. Stuart, Lisa M. Levoir, Laura Belmont, Leslie Goo

## Abstract

Antibodies targeting the so-called envelope dimer epitope (EDE) cross-neutralize Zika virus (ZIKV) and all four dengue virus (DENV) serotypes and have thus inspired an epitope-focused vaccine design against these flaviviruses. There are two EDE antibody subclasses (EDE1, EDE2) distinguished by their dependence on viral envelope (E) protein *N*-linked glycosylation at position N153 (DENV) or N154 (ZIKV) for binding. Here, we determined how E glycosylation affects *neutralization* by EDE and other broadly neutralizing antibodies. Consistent with structural studies, mutations abolishing the N153/N154 glycosylation site increased DENV and ZIKV sensitivity to neutralization by EDE1 antibodies. Surprisingly, these mutations also increased sensitivity to EDE2 antibodies although they occurred at predicted contact sites. Despite preserving the glycosylation site motif (N-X-S/T), substituting the threonine at ZIKV E residue 156 with a serine resulted in loss of glycan occupancy accompanied with increased neutralization sensitivity to EDE antibodies. For DENV, the presence of a serine instead of a threonine at E residue 155 retained glycan occupancy, but nonetheless increased sensitivity to EDE antibodies, in some cases to a similar extent as mutation at N153, which abolishes glycosylation. E glycosylation site mutations also increased ZIKV and DENV sensitivity to other broadly neutralizing antibodies, but had limited effects on ZIKV-or DENV-specific antibodies. Thus, E protein glycosylation is context-dependent and modulates the potency of broadly neutralizing antibodies in a manner not predicted by existing structures. Manipulating E protein glycosylation could be a novel strategy for engineering vaccine antigens to elicit antibodies that broadly neutralize ZIKV and DENV.

**IMPORTANCE:** Antibodies that can potently cross-neutralize Zika (ZIKV) and dengue (DENV) viruses are attractive to induce via vaccination to protect against these co-circulating flaviviruses. Structural studies have shown that viral envelope protein glycosylation is important for binding by one class of these so-called broadly neutralizing antibodies, but less is known about the determinants of neutralization. Here, we investigated how envelope protein glycosylation impacts broadly neutralizing antibody potency. By characterizing a panel of ZIKV and DENV variants encoding envelope protein glycosylation site mutations, we found that glycan occupancy was not always predicted by an intact N-X-S/T sequence motif. Moreover, envelope protein glycosylation status alters the neutralization potency of broadly neutralizing antibodies in a manner unexpected from their predicted binding mechanism as determined by existing structures. We highlight the complex role and determinants of envelope protein glycosylation that should be considered in the design of vaccine antigens to elicit broadly neutralizing antibodies.

## OBSERVATION

Zika virus (ZIKV) and the four dengue virus serotypes (DENV1-4) are closely related flaviviruses. Mature ZIKV particles are distinguished from other flaviviruses by an extended envelope (E) protein ‘150 loop’ containing a potential *N-*linked glycosylation site (PNGS) at residue N154 or N153 for ZIKV or DENV, respectively (1, 2). This PNGS is important for binding by some E dimer epitope (EDE) antibodies, which cross-neutralize DENV1-4 and ZIKV (3, 4). There are two EDE antibody subclasses, of which EDE2, but not EDE1 requires 150 loop glycosylation for efficient binding (3, 5). How glycosylation at this site impacts the potency of EDE and other broadly neutralizing antibodies (bnAbs) is less defined.

To map epitopes of antibodies that neutralize DENV1-4 but not ZIKV, we previously generated a library of DENV2 reporter virus particles (RVPs) (6) in which solvent accessible E protein residues were substituted with those corresponding to the Asian lineage ZIKV strain H/PF/2013 (7). Flavivirus RVPs have been shown to be antigenically similar to non-reporter fully infectious versions (8). The V151T mutation in the DENV2 150 loop reduced the potency of DENV1-4 bnAbs but *increased* the potency of a control EDE1 bnAb (6). To further investigate how this site impacts EDE potency, we generated RVPs encoding the reciprocal ZIKV H/PF/2013 E protein T156V mutation (**Figure S1A**) and observed up to a 32-fold increase in sensitivity to neutralization by EDE1 and EDE2 bnAbs compared to WT (**Figures 1A-D, 1H**). This mutation similarly increased sensitivity to SIgN-3C (9, 10) and F25.S02 (11) (**Figures 1E-F, 1H**), which also neutralize DENV1-4 and ZIKV but target epitopes distinct from EDE. In contrast, T156V minimally increased sensitivity (average of 3-fold IC50 reduction) to ZV-67, a ZIKV specific neutralizing monoclonal antibody (12) (**Figures 1G-H**). Compared to WT, T156V also increased sensitivity to bnAbs by at least 8-fold in the context of the ZIKV MR766 African lineage strain (13) and the Asian lineage PHL/2012 (14), but not THA/2014 (14) strain (maximum of 3-fold IC50 reduction, **Figure S2**). ZIKV THA/2014 is distinguished from other strains tested here at two E protein residues (227 and 368) (**Figure S1B**), which could have compensatory effects.

**Figure 1.**
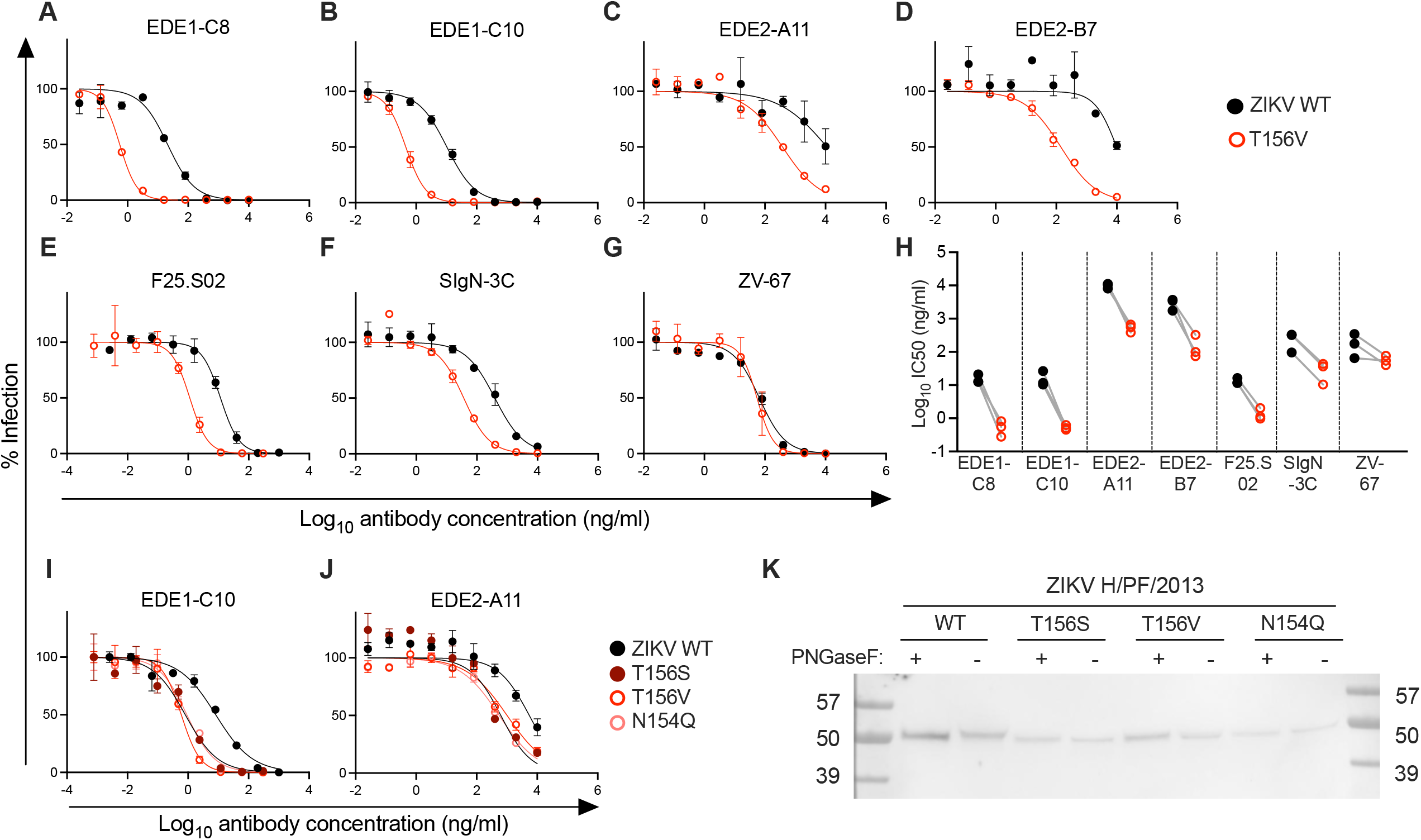
Effect of E glycosylation status on ZIKV sensitivity to bnAbs. **(A-G)** Dose-response neutralization assays using the indicated antibodies against ZIKV H/PF/2013 wild-type (WT) (black) or T156V (red) RVPs. Data are representative of three independent experiments, each performed in duplicate wells. Data points and error bars indicate the mean and range of infection in duplicate wells, respectively. **(H)** Antibody IC50 values from three independent experiments in which WT and T156V ZIKV RVPs were tested in parallel. **(I-J)** Dose-response neutralization assays using EDE1-C10 **(I)** or EDE2-A11 **(J)** against WT ZIKV H/PF/2013 RVPs or those encoding the indicated E protein mutations. Data are representative of three independent experiments, each performed in duplicate wells. Data points and error bars indicate the mean and range of infection in duplicate wells, respectively. **(K)** E proteins from untreated or PNGaseF-treated RVP lysates were detected by SDS-PAGE and Western blot. Size markers (kDa) are shown in leftmost and rightmost lanes. Data are representative of seven independent experiments performed using two independent WT and mutant RVP stocks prepared in parallel.

The T156V mutation abrogates the only ZIKV E protein PNGS (**Figure S1A**) (1, 15). Increased ZIKV sensitivity to bnAbs conferred by this mutation is in contrast to the previous observation that abolishing this PNGS did not impact sensitivity to poorly neutralizing antibodies targeting the E protein fusion loop (16). To determine if increased sensitivity of ZIKV T156V to bnAbs is specifically due to loss of the PNGS, we generated ZIKV H/PF/2013 RVP variants containing additional mutations at E residues N154 and T156. Each of these variants, including those that abolish the PNGS, efficiently infected Raji-DCSIGNR cells (**Figure S3A**) even though cellular attachment depends on interactions between DCSIGNR and viral glycans (17). As previously suggested, the presence of glycosylated, uncleaved prM retained on the virion surface due to incomplete maturation likely facilitates attachment in the absence of the E PNGS (16). Accordingly, ZIKV RVPs encoding E protein PNGS mutations prepared in the presence of exogenous furin, which improves prM cleavage efficiency, displayed a markedly reduced ability to infect Raji-DCSIGNR cells compared to corresponding virus stocks prepared using standard methods (**Figures S3D-E**). In contrast, WT ZIKV prepared with or without excess furin displayed relatively similar infectivity (**Figure S3B**). Despite retaining the PNGS motif (N-X-S/T), the infectivity of T156S RVPs prepared with or without furin resembled that of variants that ablate this motif (**Figure S3C**), suggesting loss of E glycosylation.

Like the T156V mutation, ZIKV encoding a N154Q or N154A mutation that disrupts the PNGS was more sensitive to neutralization by EDE1-C10 (**Figure 1I**) and EDE2-A11 than WT ZIKV (**Figure 1J**). For EDE1-C10, this finding is consistent with structural studies suggesting that EDE1 antibodies displace the glycan-containing 150 loop to interact with the E protein (5). Surprisingly, despite an intact PNGS motif, ZIKV T156S was also more sensitive to EDE1-C10 and, to a lesser extent, EDE2-A11, compared to WT (**Figures 1I-J**). SDS-PAGE and western blotting of RVP lysate confirmed that WT ZIKV E protein displayed a slightly lower molecular weight after PNGaseF treatment, demonstrating glycan cleavage (**Figure 1K**). In contrast, ZIKV T156S E protein migration was unaffected by PNGaseF treatment, similar to E protein from ZIKV N154Q or T156V, each of which ablates the PNGS motif. Combined with its infectivity (**Figure S3C**) and neutralization profiles (**Figure 1I-J**), this finding indicates that the E protein of ZIKV T156S produced in mammalian cells was not glycosylated. The relative inefficiency of *N*-linked glycosylation associated with the presence of a serine versus a threonine within the PNGS motif (18) has been similarly described for a rabies virus glycoprotein (19).

To investigate how glycosylation of the DENV 150 loop impacts neutralization sensitivity, we generated DENV2 16681 RVP variants encoding mutations at E residues N153 and T155. Unlike our findings with ZIKV, the migration pattern of E protein of DENV2 T155S with or without PNGaseF treatment was similar to that of WT DENV2, demonstrating intact glycosylation (**Figure 2H**). As expected, E protein from DENV2 encoding the N153Q or T155A mutation, each of which abrogates the PNGS at this position, migrated faster than WT or T155S E protein, indicating loss of glycan occupancy.

**Figure 2.**
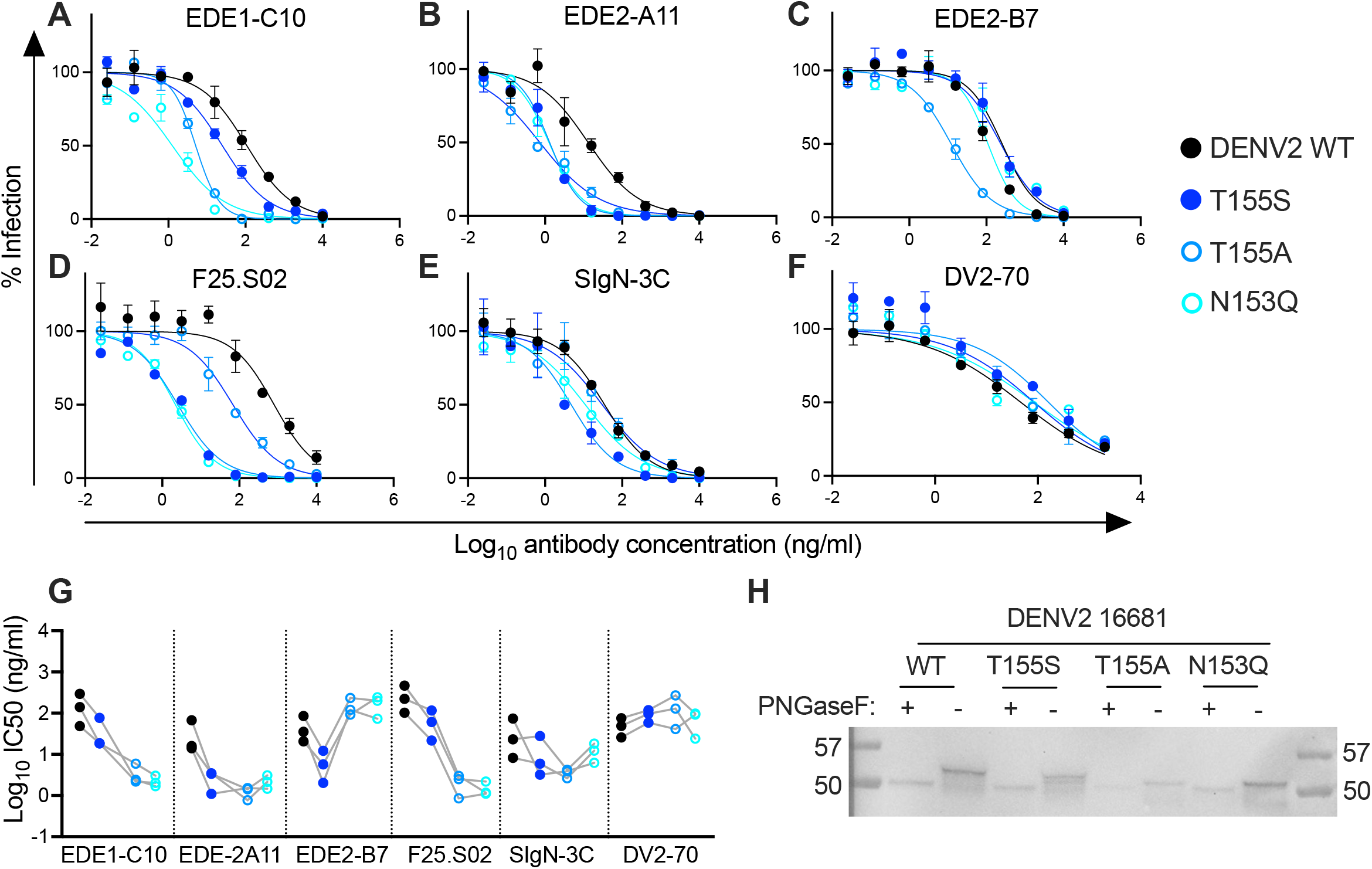
Effect of E glycosylation status on DENV2 sensitivity to bnAbs. **(A-F)** Dose-response neutralization assays using the indicated antibodies against WT DENV2 16681 RVPs or those encoding the indicated E protein mutation. Data are representative of three independent experiments, each performed in duplicate wells. Data points and error bars indicate mean and range of infection in duplicate wells, respectively. **(G)** Antibody IC50 values from three independent experiments using antibodies shown on the x-axis. In each experiment, WT and DENV2 16681 RVPs encoding the E protein mutations shown in A-F were tested in parallel. **(H)** E proteins from untreated or PNGaseF-treated RVP lysates were detected by SDS-PAGE and Western blot. Size markers (kDa) are shown in leftmost and rightmost lanes. Data are representative of six independent experiments performed with two independent WT and mutant RVP stocks prepared in parallel. DENV E protein has two PNGS motifs at residues N67 and N153 (Figure S1A); only mutations that affect the latter are assessed here. The multiple bands observed in untreated wells likely indicate glycosylation at either one or both PNGS.

Although DENV2 mutations that do (T155S) or do not (T155A, N153Q) maintain the 150 loop PNGS motif minimally (<3-fold) impacted the potency of DV2-70, a DENV2-specific antibody (20) (**Figures 2F-G**), each increased bnAb potency to various extents. The largest effect seen for bnAb SIgN-3C was against T155S RVPs (~10-fold increase in sensitivity, **Figures 2F-H**). In contrast, compared to T155A or N153Q (100-to 300-fold increase in sensitivity), the T155S mutation increased sensitivity to EDE1-C10 and F25.S02 relatively modestly (4-to 10-fold) (**Figures 2A, D, G**). All three mutations similarly increased sensitivity to EDE2-A11 neutralization by ~20-fold (**Figures 2B, G**). Thus, in addition to E glycosylation status, the specific amino acid chemistry within the PNGS motif modulates DENV2 sensitivity to bnAbs.

The observed increased neutralization potency associated with the loss of the DENV2 150 loop PNGS was unexpected especially for EDE2-A11, which is predicted to contact the N153 glycan based on structural studies with soluble E protein (5). Mutagenesis of DENV subviral particles also showed that this PNGS was important for EDE2 binding (3). When tested against EDE2-B7, a somatic variant of EDE2-A11 (5), loss of PNGS via N153Q or T155A mutation rendered DENV2 less sensitive to neutralization (**Figures 2C, G**), which is consistent with the above prior structural studies. Nevertheless, the T155S mutation, which preserves glycosylation, *increased* DENV2 sensitivity to EDE2-B7 by 124-fold (**Figure 2C, G**). These findings demonstrate that DENV2 E glycosylation has distinct impacts even on bnAbs with highly similar structural epitopes and genetic characteristics.

### Conclusions

Flavivirus E protein glycosylation is context-dependent, and variation at the PNGS within the 150 loop modulates sensitivity to bnAbs in ways not predicted by existing structural studies. We underscore the complex determinants and antigenic consequences of E glycosylation; both are impacted by even a conservative change within the PNGS motif. Notably, there is variation at the PNGS within the 150 loop of currently circulating DENV strains (21). Our findings suggest that mutations within this PNGS are antigenically relevant and should be considered in the design of vaccine antigens to elicit bnAbs.

## METHODS

### Cells

Raji-DCSIGNR cells (a gift from Ted Pierson, NIH) were maintained in RPMI 1640 medium supplemented with GlutaMAX (Cat# 72400–047; ThermoFisher Scientific), 7% Fetal bovine serum (FBS)(Cat# 26140079; ThermoFisher Scientific), and 100 U/mL penicillin-streptomycin (Cat# 15140122, ThermoFisher Scientific).

HEK-293T/17 cells (Cat# CRL-11268, ATCC) were maintained in DMEM (Cat# 11965118; ThermoFisher Scientific) supplemented with 7% FBS and 100 U/mL penicillin-streptomycin. Raji-DCSIGNR and HEK293T/17 cells were generally maintained at 37° C with 5% CO2. However, for RVP production, HEK-293T/17 cells were maintained at 30° C in a low-glucose formulation of DMEM (Cat # 12320–032; ThermoFisher Scientific), as previously described (22).

Expi-CHO-S cells (Cat# A29127; ThermoFisher Scientific) were cultured in ExpiCHO Expression Medium (Cat# A2910001; ThermoFisher Scientific) and maintained at 37°C in 8% CO2 on a platform rotating at 125 rpm with a rotational diameter of 19 cm.

### Generation of plasmids encoding E protein variants

Previously described plasmids encoding the structural genes (C-prM-E) of DENV2 16681, ZIKV H/PF/2013, ZIKV MR766, ZIKV THA/2014, and ZIKV PHL/2012 (8, 23) (all provided by Ted Pierson, NIH) were used as templates for Q5 site-directed mutagenesis (Cat# E0554S; New England Biolabs,) using primers designed with NEBaseChanger (New England Biolabs). Whole plasmids were confirmed by Oxford Nanopore sequencing (Plasmidsaurus) to confirm only the desired mutation was present.

### Production of reporter virus particles (RVPs)

HEK-293T/17 cells were co-transfected with 1 µg of a plasmid expressing a West Nile virus subgenomic replicon in which the structural genes have been replaced with GFP (24), and 3 µg of a plasmid expressing the WT or mutant C-prM-E structural genes of interest using Lipofectamine 3000 (Cat# L3000-015; ThermoFisher Scientific) according to manufacturer’s protocol. After 4 hours of incubation at 37°C, media was replaced and cells transferred to 30°C. Virus-containing supernatant was collected on days 3-6 post-transfection, pooled, passed through a 0.22 µm Steriflip filter (Cat# SE1M179M6, Millipore-Sigma), aliquoted, and stored at -80°C. Mature RVP stocks were produced using the same protocol, except with the addition of 1 µg of a human furin-expressing plasmid (provided by Ted Pierson, NIH).

### Determining infectious titer of RVPs

Raji-DCSIGNR cells (2e5/well in 20 µl) were infected with 10 × 2-fold serial dilutions of an equal volume of each RVP stock in 384-well plates (Cat# 164688, ThermoFisher Scientific) and incubated at 37°C. Two days later, cells were fixed in a final concentration of 2% paraformaldehyde (Cat# 15714S; Electron Microscopy Sciences) and % GFP positive cells were enumerated by flow cytometry (Intellicyt iQue Screener PLUS, Sartorius AG). Infectious titers were determined from the most linear portions of the dose-response infectivity curves using the following formula: (((%GFP positive cells/100) * number of cells infected) / volume of virus) * virus dilution factor.

### Production and purification of antibodies

Heavy and light variable region sequences of F25.S02 were obtained by single B cell transcriptomics as previously described (11). Variable region sequences for other bnAbs were determined based on the following protein database (PDB) IDs: 4UT9 (EDE1-C10), 4UTA (EDE1-C8), 4UT6 (EDE2-B7), 4UTB (EDE2-A11), and 7BUD (SIgN-3C). Corresponding nucleotide sequences were codon-optimized, synthesized (Twist Bioscience, South San Francisco, CA), and cloned into mammalian expression vectors provided by Patrick Wilson (University of Chicago): AbVec-hIgG1 (GenBank accession # FJ475055) and AbVec-hIgKappa (GenBank accession# FJ475056) or AbVec-hIgLambda (GenBank accession # FJ517647). All AbVec antibody expression plasmids (IgG1-heavy, kappa, and lambda) were confirmed by Sanger sequencing using the primer “AbVec sense”: GCTTCGTTAGAACGCGGCTAC. Sequence-confirmed heavy and light chain plasmids were co-transfected into cultures of ExpiCHO-S cells at 0.8 ng/mL total DNA concentration at 1:1 mass ratio using OptiPro serum free medium (Cat#12309, Gibco) and Expifectamine CHO Transfection Kit (Cat# A29130, Gibco) according to the manufacturer’s instructions. Eight days post-transfection, IgG-containing supernatant was collected, clarified by centrifugation (3220 x g for 10 minutes), and filtered through a 0.45 µm membrane. IgG was purified using MabSelect Sure LC protein A agarose beads (Cat# 17-5474-01, Cytiva Life Sciences) according to the manufacturer’s instructions. Mouse monoclonal antibody 4G2 was purified from the hybridoma D1-4G2-4-15 (Cat# HB-112, ATCC) by the Fred Hutchinson Cancer Center Antibody Technology Core.

### Neutralization assays

Neutralization assays were performed in 384-well plates. RVPs were diluted to 5-10% infectivity (20 µl/well) and incubated for 1 hour at room temperature with 10 × 5-fold serial dilutions of an equal volume of antibodies before infection of an equal volume of Raji-DCSIGNR cells (2e5/well) at 37° C for 2 days. Cells were fixed with 2% paraformaldehyde and % GFP positive cells were determined using flow cytometry (Intellicyt iQue Screener PLUS, Sartorius AG). Infection was normalized to conditions without antibody. The antibody concentration that neutralized 50% infection (IC50) was estimated by non-linear regression with a variable slope and the bottom and top of the curves constrained to 0% and 100%, respectively (GraphPad Prism 9.5.1).

### Determining E protein glycosylation status

RVP stocks prepared as described above were concentrated by microcentrifugation overnight (20,000 xg at 4°C) through a 20% sucrose cushion (0.25 ml sucrose per 1 ml supernatant). RVP pellets were resuspended in HNE buffer (5mM HEPES, 150mM NaCl, 0.1mM EDTA, pH adjusted to 7.4) For SDS-PAGE, concentrated RVPs were lysed by incubating at 55°C for 20 min with Bolt LDS sample buffer (Cat# B0007, ThermoFisher Scientific). E protein glycosylation status was determined by treating RVP lysates with PNGase F (Cat#P0704S, New England Biolabs) for 3 hours at 37°C. Treated and untreated RVP lysates were run on a 4-12% BisTris gel (Cat# NW04120BOX, ThermoFisher Scientific) for 1 hour at 100V, and transferred to a nitrocellulose membrane using the Invitrogen iBlot system (Cat# IB21001, ThermoFisher Scientific).

Membranes were blocked for 1 hour at room temperature in 3% milk diluted in 1X Tris-buffered saline and Tween 20 (TBS-T), followed by overnight incubation with rocking at 4° C with 3 µg/ml mouse primary anti-E antibody, 4G2 (purified from hybridoma as described above), or ZV-67 (a gift from Michael Diamond, Washington University, St. Louis) to detect E protein of DENV2 or ZIKV, respectively. The next day, membranes were washed three times with TBS-T, followed by incubation for 1 hour at room temperature with a horseradish peroxidase conjugated anti-mouse secondary antibody (Cat# NA931-1ML, Millipore Sigma) diluted 1:1000 in 3% milk. Membranes were again washed three times with TBS-T and protein bands detected using chemiluminescence (Cat# RPN3004, Cytiva Life Sciences).

## ACKNOWLEDGEMENTS

We thank Ted Pierson for providing Raji-DCSIGNR cells and constructs for generating reporter virus particles; Michael Diamond for providing mouse monoclonal antibodies, ZV-67 and DV2-70; Patrick Wilson for providing IgG expression vectors; and members of the Goo lab for helpful discussion. This work was supported by the Fred Hutchinson Cancer Center Diverse Trainee Fund (MC); Diseases of Public Health Importance Training Grant 2T32AI007509 (JBS); Viral Pathogenesis and Evolution Training Grant T32 AI083203 (LB); and the Antibody Technology (RRID:SCR_022608) Shared Resource Facilities of the Fred Hutch/University of Washington/Seattle Children’s Cancer Consortium (P30 CA015704).

