## Supplementary Figure 1 for "Defining the impact of flavivirus envelope protein glycosylation on sensitivity to broadly neutralizing antibodies"

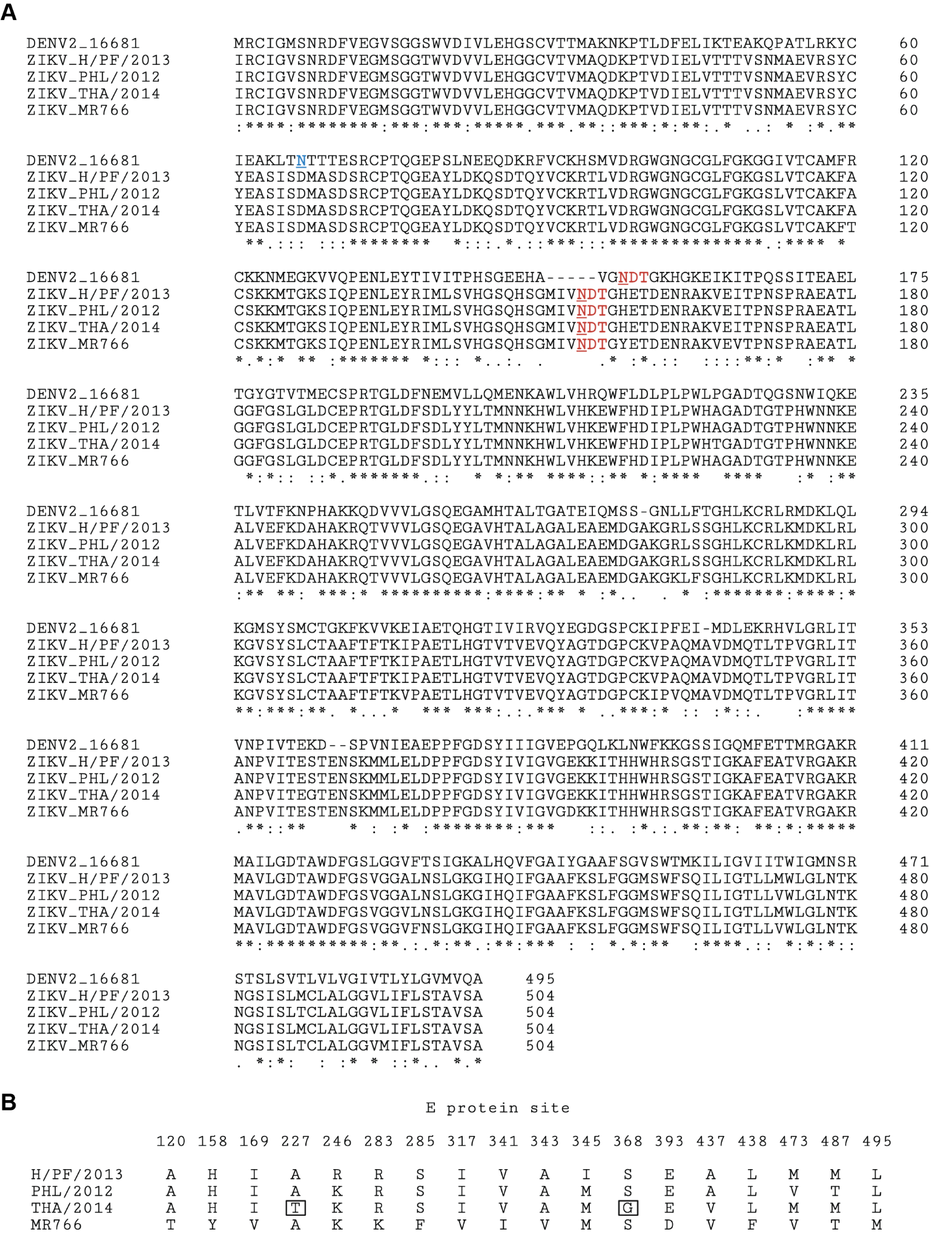

**Figure S1. E protein sequence alignment of ZIKV and DENV. (A)** Multiple sequence alignment of the indicated DENV2 and ZIKV E protein sequences created using Clustal Omega tool (EMBL-EBI) (25). Genbank Genome Accession numbers are as follows: NC_001474.2 (DENV2_16681), KJ776791.2 (ZIKV_H/PF/2013), KU681082.3 (ZIKV_PHL/2012), KU681081.3 (ZIKV_THA/2014), HQ234498.1 (ZIKV_MR766). The PNGS motif in the 150 loop is shown in red. Within this motif, site N153 or N154 in DENV2 or ZIKV, respectively, is underlined. The additional PNGS at DENV2 residue N67 is shown in blue. Under the alignment, ‘*’ indicates completely conserved residue; ‘:’ indicates amino acids with strongly similar properties (score of >0.5 in the Gonnet PAM 250 matrix) and ‘.’ indicates amino acids with weakly similar properties (score of <=0.5 in the Gonnet PAM 250 matrix). **(B)** E protein sites of amino acid variation among the ZIKV strains tested in this study (Genbank Genome Accession numbers as in (A)). E protein sites and amino acid residues that distinguish ZIKV strain THA/2014 from the others are boxed.
