## Supplementary Figure 2 for "Defining the impact of flavivirus envelope protein glycosylation on sensitivity to broadly neutralizing antibodies"

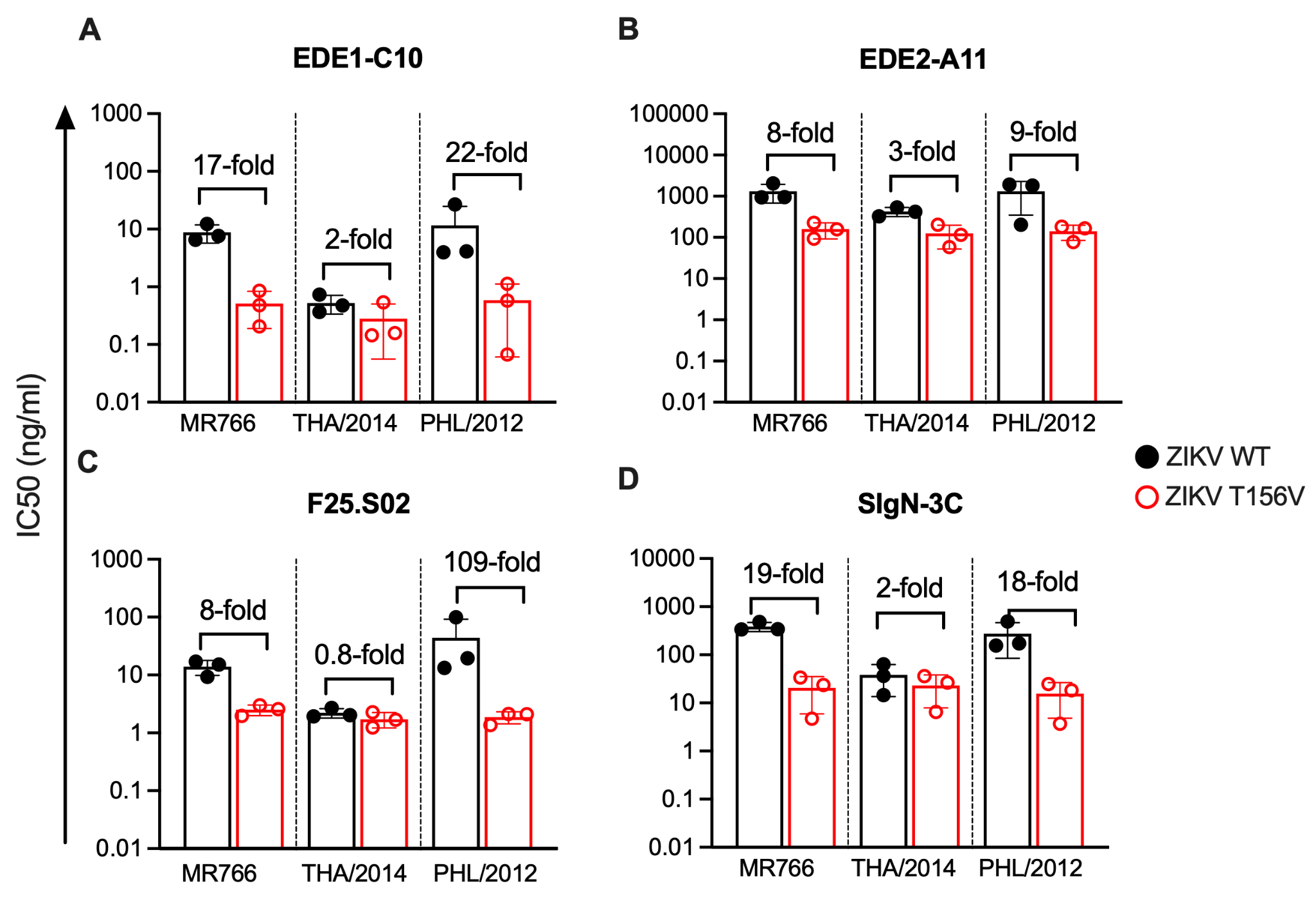

**Figure S2. Effect of T156V mutation on sensitivity of additional ZIKV strains to neutralizing antibodies**

**(A-E)** Bar graphs show mean IC50 values of the indicated antibodies against WT (black) or T156V (red) RVPs of ZIKV strains (x-axis) from three independent experiments, each represented by a data point. Error bars indicate the standard deviation.
