## Supplementary Figure 3 for "Defining the impact of flavivirus envelope protein glycosylation on sensitivity to broadly neutralizing antibodies"

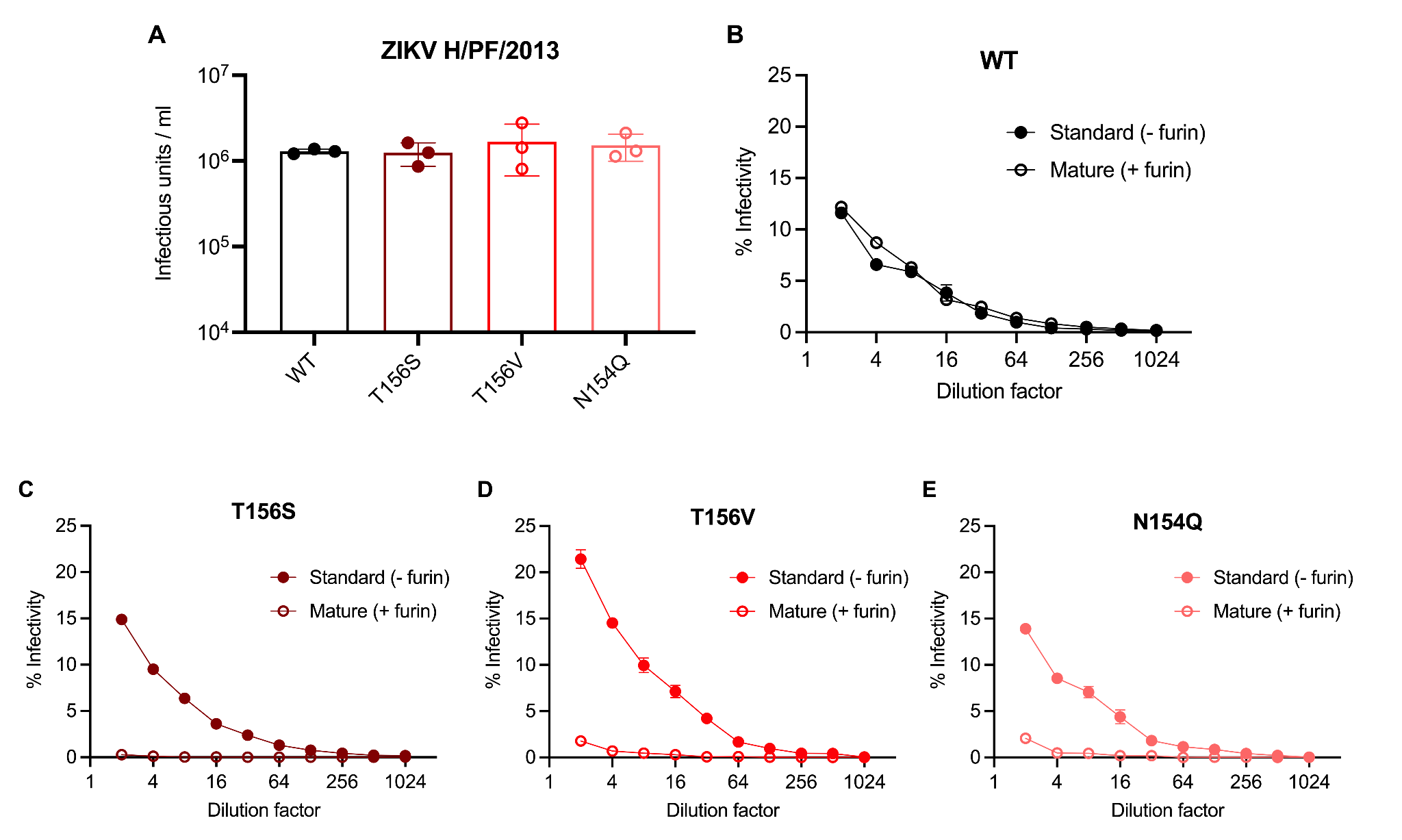

**Figure S3. Infectious titer of ZIKV H/PF/2013 RVPs on Raji-DCSIGNR cells**

**(A)** Bar graphs show the mean infectious titers from three experiments (data points) performed using three independent standard preparations of RVPs. Error bars indicate the standard deviation. **(B-E)** Dose-response infectivity curves of the indicated ‘standard’ or ‘mature’ ZIKV H/PF/2013 RVPs prepared in the absence (filled circles) or presence (open circles) of exogenous furin, respectively. Data shown is from one experiment performed in duplicate wells; error bars indicate the range of infection.
